# MycoMobilome: A community-focused non-redundant database of transposable element consensus sequences for the fungal kingdom

**DOI:** 10.1101/2025.10.28.685023

**Authors:** Tobias Baril, Daniel Croll

**Affiliations:** Laboratory of Evolutionary Genetics, Institute of Biology, University of Neuchâtel, Rue Émile-Argand 11, 2000 Neuchâtel, Switzerland

## Abstract

Transposable elements (TEs) are found in nearly all eukaryotic genomes. Despite significant advances in the sequencing of genomes, TE resources remain sparse, leading to a lack of traceability, reproducibility, and duplication of effort when annotating TEs. Here, we focus on the fungal kingdom and present MycoMobilome, a non-redundant database of TE consensus sequences systematically curated using a set of 4,309 genomes covering all major clades. The initial database contains 276,641 consensus sequences after filtering to remove putative host genes and low-quality consensus sequences. We provide a consistent naming convention to surface information on the confidence in the classification including potential conflicting ORF functions, along with metadata to enable evaluation of TEs of interest and to determine whether further curation work is required on a case-by-case basis. Finally, we provide guidelines for community contributions, and encourage researchers to deposit new or curated sequences, which will be incorporated into future MycoMobilome releases.

## Introduction

Annotation of transposable elements (TEs) is essential for the analysis of genomes even if the focus is primarily on coding sequences. TE annotation relies on the initial discovery of TE families, termed *de novo* curation, followed by homology-based annotation using tools to recognise all members of each TE family. Various approaches exist for *de novo* curation, with the most common including RepeatModeler2 (1) and TEdenovo of REPET (2), whilst there are also databases providing previously curated TE families that can be used to annotate genomes, including Dfam (3, 4), TREP (5), and RepBase (6).

The fungal kingdom spans almost a billion years of evolution, exhibits remarkable phenotypic and genomic diversity, and contains species found in nearly all ecological niches. The availability of genome resources for species spanning the kingdom is accelerating, partly due to efforts including the 1000 fungal genomes initiative (https://1000.fungalgenomes.org) and broader efforts of the Darwin Tree of Life and Earth Biogenome projects (7, 8). Despite these growing efforts, curated and multi-species TE resources for the fungal kingdom are almost non-existent. For example, in Dfam version 3.9, the only curated TEs are found for *Septoria linicola* and *Zymoseptoria tritici* (https://dfam.org/). Dedicated efforts to curate TE libraries for individual species include the plant pathogens *Magnaporthe oryzae* (9) and *Z. tritici* (10). However, cross-referencing TE annotations across species has not been attempted, leading to reproducibility issues and duplication of effort as researchers perform annotations largely on a per-species basis.

Here, we present MycoMobilome (https://github.com/TobyBaril/MycoMobilome), a non-redundant database of TE consensus sequences covering species across the fungal kingdom. We created MycoMobilome using all publicly available genome resources for fungi and provide researchers with a systematically generated TE consensus library with a persistent naming scheme to improve consistency and reproducibility. Consensus sequences are provided with labels to show whether their classification is supported by open reading frames (ORFs), and whether these consistently match the provided classification. Applying MycoMobilome classification to primary fungal models, we find that the genome fraction annotated as TEs increases by between +7.20% in *M. oryzae* to +13.41% in *Cryptococcus neoformans* compared to existing community TE annotations.

MycoMobilome is a key resource to facilitate investigations into TEs across the fungal kingdom and provides easy-to-implement methods for researchers to contribute new or updated sequences with attribution. Consistent application of MycoMobilome will improve consistency in TE naming conventions, whilst reducing the duplication of effort often accompanying repeat annotation in new genome assemblies.

## Materials and Methods

### Initial Curation for MycoMobilome

All publicly available genomes and associated proteomes for the fungal kingdom were sourced using Mycotools (version 0.32.3)(11). Genomes were manually filtered to remove those under embargo, resulting in a final set of 4,309 genome assemblies. Genome assembly metadata is provided in the MycoMobilome database (https://doi.org/10.5281/zenodo.17037469).

For each genome, a library of putative TE consensus sequences was generated using `earlGreyLibconstruct` in Earl Grey (v4.4.0)(12), configured with Dfam curated elements only (v3.7)(3, 4). All 4,309 *de novo* consensus libraries were subsequently combined into a single FASTA file containing 773,843 sequences. These sequences were clustered using the `easy-cluster` scalable cascaded clustering approach in MMseqs2 (13) with `--min-seq-id 0.8 -c 0.8 --cov-mode 1 --cluster-reassign` to cluster sequences to the 80-80-80 TE family rule (14), resulting in 354,315 non-redundant consensus sequences. The representative sequence for each cluster was extracted and labelled with the source genome (*i.e*. the genome from which the consensus sequence originated).

Autonomous TEs encode domains for their selfish activity, and the identity of these open reading frames (ORFs) can be used to classify TEs (14–16). Conversely, multicopy host proteins can erroneously be curated as repetitive sequences by automated TE curation methods, given their repetitive occurrence in host genomes. We translated all six frames of each TE consensus sequence using `transeq -clean -frame 6` in EMBOSS (v.6.6.0)(17). We identified matches to known host proteins present in the Fungi partition of RefSeq release 228 (18) using Diamond BLASTp (v 2.1.11) (19) with `--sensitive --matrix BLOSUM62 --evalue 1e-3`. Following this, we detected similarity to characterised TE proteins using two complementary approaches. First, we used HMMscan in HMMER (v3.4)(20) with `-E 10 --noali` to detect homology to known TE protein domain hmm models supplied in ProfilesBankForREPET_Pfam35.0_GypsyDB as part of the REPET software suite (2, 21, 22). Hits were filtered to retain those with fseq_evalue ≤ 0.001 and fseq_bitscore ≥ 50. Next, we used BLASTp (v2.14.1+)(23) to detect homology to known TE protein domains found in RepeatPeps.lib, which is part of RepeatMasker (v4.1.9)(24) and is used to classify TEs using the RepeatClassifier module in the initial *de novo* curation step in Earl Grey. A minimum e-value of 1e-3 was defined, and nested hits were removed to retain the highest quality protein hit for each query, followed by combining of adjacent and overlapping hits. Potential host gene hits were identified using the approach developed by (25). A TE consensus sequence was designated as a putative host gene if either: (i) there were hits to RefSeq queries and no hits to known TE queries; (ii) there were hits to both RefSeq queries and known TE queries, but at least 90 residues aligned to a RefSeq query did not overlap with alignments to known TE queries. Consensus sequences with no hits were retained in the TE library as putative non-autonomous TEs. In total, 24,571 consensus sequences were identified as potential host genes and removed.

To further refine the database, poor-quality TE consensus sequences were filtered. We define a poor-quality consensus sequence as <120bp in length, as these are likely to be incomplete and poor quality. Further, majority-rule consensus sequences can sometimes contain unknown nucleotides, indicated with N in the sequence. We removed all sequences with ≥5% N using `seqtk comp` (https://github.com/lh3/seqtk). In total, 53,103 consensus sequences were removed for not meeting the quality thresholds, resulting in a final database size of 276,642 putative TE consensus sequences.

The majority of TE consensus sequences in publicly available databases remain uncurated. Despite this, these sequences remain a widely used resource. To enable the community to evaluate TE annotation quality of newly analyzed genomes, we introduce an automated classification of each TE consensus against protein hits to known TEs (Tables are provided in the MycoMobilome database). Each TE consensus was labelled with a two-letter code providing the level of evidence supporting the TE classification: `_PE` for protein evidence that matches the given classification, `_DA` for protein evidence that contradicts the given classification, and `_NE` for no protein evidence, which includes non-autonomous TEs.

Finally, each TE consensus in MycoMobilome was given a unique name with the following format: `MycMob1.0_family-[unique_family_number]- [six_letter_species_code]_[protein evidence]#[high_level_TE_classification]/[sub_level_TE_classification] @[genus species]`.

### Comparison of existing TE annotation approaches with MycoMobilome

We wanted to assess the extent to which sampling TEs across fungal diversity enables improved detection of TE-derived genome content. To do this, we selected three well- studied organisms and compared TE annotation using MycoMobilome against the traditional approaches employed by the research communities working with each of these organisms. We compared our standardised approach with MycoMobilome against (i) annotation with known fungal TEs from RepBase and Dfam, as used in a recent study on *C. neoformans* (26); (ii) a community annotation resource for *Candida albicans* (http://www.candidagenome.org/download/gff/C_albicans_SC5314/Assembly22/); (iii) annotation using a curated TE library with RepeatMasker for *M. oryzae* (9).

For *C. neoformans*, we adopted the same approach used in the original study and annotated the genome assembly for isolate CNA3 H99 (GCF_000149245.1) using RepeatMasker (v4.1.9) configured with Dfam (v3.9) and RepBase RepeatMasker Edition (release 20181026) and the options `-species fungi -norna -no_is`. For *M. oryzae*, we sourced the reference genome assembly MG8 (GCF_000002495.2) and annotated this with RepeatMasker using the TE consensus library provided with the publication (9). For *C. albicans*, we obtained the genome assembly in FASTA format and the community feature annotation in GFF format for genome assembly SC5314 Assembly 22 from the Candida genome portal (http://www.candidagenome.org/download/gff/C_albicans_SC5314/Assembly22/). To obtain a GFF of repeat regions, we filtered the feature GFF for features named long_terminal_repeat, repeat_region, and retrotransposon.

For comparison, we adopted our standardised approach as recommended with MycoMobilome. We annotated all three genome assemblies with `earlGreyAnnotationOnly`using Earl Grey (v6.3.2), providing MycoMobilome as the input library, with all other settings left as default.

Hits that are shared and unique to each methodological approach were identified using BEDTools intersect (v2.31.1)(27), and resultant feature coverage was calculated in R (v4.4.3)(28) with tidyverse (29). For shared annotations, the width of the annotation in each case was calculated, rather than using a single length of annotation across compared methodologies, to account for variation in annotation size between methods. Venn diagrams were generated using the ggVennDiagram package (30).

## Results & Discussion

### MycoMobilome assesses TE evidence across the fungal kingdom

The MycoMobilome release contains 276,642 consensus sequences, of which 39,265 have classifications supported by protein evidence. This provides a curated, non-redundant resource for the diversity of TEs across the fungal tree of life. In particular for clades without previous TE annotation efforts, this represents a very substantial expansion of genomic datasets. We performed filtering steps to stringently remove any consensus sequences with the potential to be host genes, as well as poor-quality consensus sequences. Following these efforts, we provide three versions of the database: a full database, a subset containing only TE consensus sequences with protein evidence, and a subset containing only TE consensus sequences lacking protein evidence. The compressed database files are provided via Zenodo (https://doi.org/10.5281/zenodo.17037469) in FASTA format, ensuring compatibility with a broad range of bioinformatics approaches. We provide guidance to non-specialists on how to best implement the MycoMobilome dataset into TE annotation pipelines (https://github.com/TobyBaril/MycoMobilome). We also provide a record of the genomes used to curate TEs across fungal diversity, and tables of hits to known TE proteins from multiple sources to cross-reference evidence supporting TE classifications for sequences of interest.

### A community-based curation effort through continuous submissions

We also provide a MycoMobilome community (https://zenodo.org/communities/mycomobilome), hosted on Zenodo, with the aim of encouraging users to contribute new or improved (*i.e*. manually curated) TE consensus sequences to future releases. TE curation takes considerable work and expertise, which researchers should be recognised for. By adopting the Zenodo community approach, user contributions are assigned persistent DOIs, enabling contributors to be cited and recognised for their contributions to the wider genomics community. We provide guidance on how to contribute to MycoMobilome on the GitHub page, maximising ease-of-use. Further, hosting such a resource on Zenodo reduces risks associated with longevity of the database. The nature of MycoMobilome as a database in FASTA format limits file sizes and enables efficient storage and reuse without extensive infrastructure or resource requirements.

### MycoMobilome improves TE detection in important fungal models

By sampling TEs across fungal diversity, we increase the proportion of well-studied host genomes that is attributed to TEs (Figure 1, Supplementary Table 1). For the major rice and wheat pathogen, *M. oryzae*, which has a well-curated TE library, and much attention focused on TE dynamics, MycoMobilome increases the proportion of the host genome annotated as TEs by 7.20% (2.95Mb). The use of Earl Grey to annotate TEs also increases the proportion of the genome annotated as TEs for loci detected both in previous annotations and MycoMobilome by 10.57% (*i.e*. “shared”; 801.1kb). The improved annotation is likely due to the incorporation of defragmentation and overlap resolution steps, consistent with previous observations (12). When annotating the genome of *C. neoformans* with MycoMobilome, we find an increase of host genome proportion annotated as TEs of 13.41% (2.54Mb) compared to the previous annotation, which relied on fungal TEs hosted in Dfam and RepBase (Figure 1). In addition, a similar pattern is observed in *C. albicans*, where annotation using MycoMobilome increases the proportion of the genome recognised as TE by 9.82% (2.81Mb) compared to the current community resource based on within-species TE characterisation. Hence, MycoMobilome provides a substantial increase in well-characterized TEs even in comparatively well-studied fungal models.

**Figure 1.**
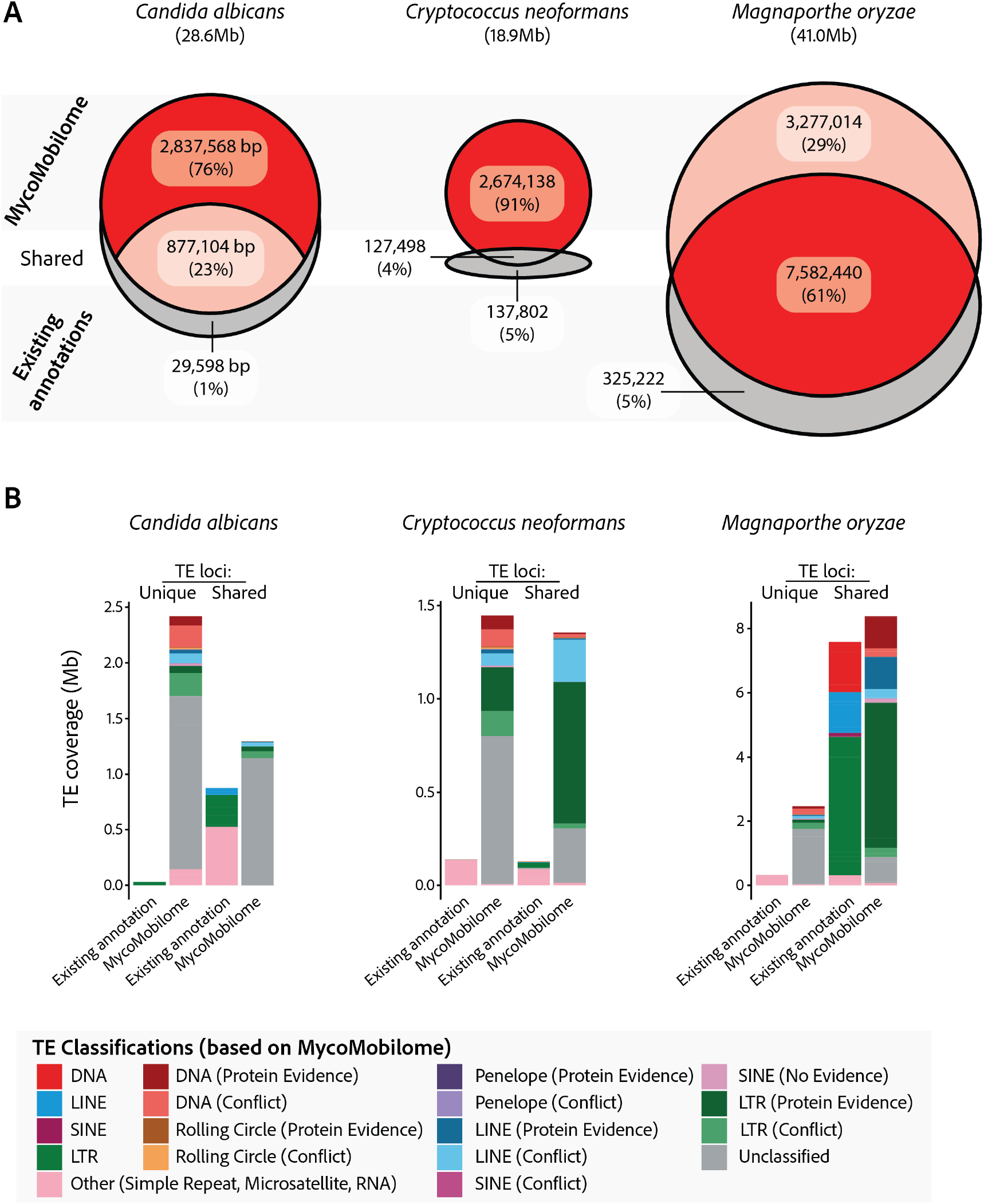
The use of MycoMobilome increases the fraction of genomes annotated as transposable elements in three key fungal pathogen models. **A.** Venn diagrams illustrating the number of base pairs for each genome assembly that were annotated as TEs using either classical approaches or MycoMobilome (with Earl Grey). Depth of colour and plot area is proportional to the percentage of total base pairs annotated as TE across all categories. Numbers in brackets show the percentage of all annotated base pairs that are found in each category, as labelled. Area of Venn diagrams is proportional to total genome size. **B**. TE annotations split by approaches with which they are identified, and the classification of each annotated TE, as indicated in the key. Discrepancies in TE coverage at shared loci arise due to the defragmentation and overlap resolution steps automatically performed with Earl Grey following TE annotation with MycoMobilome.

TEs can persist in host genomes over long evolutionary time, over which they will experience mutation and degradation leading to the accumulation of genomic fossils, challenging our ability to detect these sequences as TE-derived. However, in related lineages, these TE families may persist and be retained with higher levels of identity and at higher copy numbers, enabling their detection. By sampling TEs across deep evolutionary time and using this information to generate a non-redundant TE library, we were able to detect homology to TEs that may evade detection using single-species *de novo* approaches, which require high TE copy numbers to generate an initial consensus sequence. We show that the use of MycoMobilome increases the proportion of the host genome annotated as TEs, demonstrating the power of sampling TEs across the fungal kingdom, and over deep evolutionary time, to be able to detect TE-derived sequences across fungal diversity.

## Conclusions

Here, we introduce MycoMobilome, a non-redundant database of TE consensus sequences for the fungal kingdom. We aim to provide this database as a key starting point to facilitate genomic investigations in lineages spanning the fungal kingdom. Key features include consistent naming conventions, metadata on the quality of TE classifications and an easy-to-follow tutorial for the community to process additional genomes. We welcome contributions to the MycoMobiliome database and provide citable recognition for submitted TE annotations. We encourage the fungal research community to engage and expand the database scope to the benefit of all.

## Data Availability

The MycoMobilome database has been deposited in the Zenodo database under DOI 10.5281/zenodo.17037468 (https://doi.org/10.5281/zenodo.17037468). User guidance and documentation is hosted on GitHub (https://github.com/TobyBaril/MycoMobilome) and under the DOI 10.5281/zenodo.17473060 (https://doi.org/10.5281/zenodo.17473060).

## Acknowledgements

DC was supported by the Swiss National Science Foundation (grant 201149) and by an Innosuisse grant (32532.1 IP-LS).

## Author Contributions Statement

T.B. and D.C. conceived and coordinated the study. T.B. performed analyses. D.C. provided funding and supervised the work. T.B and D.C. wrote the manuscript.

